# mTORC1 Activation In Presumed Classical Monocytes: Observed Correlates With Human Size Variation and Neuropsychiatric Disease

**DOI:** 10.1101/2024.01.31.578140

**Authors:** Karl Berner, Naci Oz, Alaattin Kaya, Animesh Acharjee, Jon Berner

## Abstract

**Background:** Gain of function disturbances in nutrient sensing are likely the largest component in human age-related disease. Mammalian target of rapamycin complex 1 (mTORC1) activity affects health span and longevity. The drugs ketamine and rapamycin are effective against chronic pain and depression, and both affect mTORC1 activity. Our objective was to measure phosphorylated p70S6K, a marker for mTORC1 activity, in individuals with psychiatric disease to determine whether phosphorylated p70S6K could predict medication response.

**Methods:** Twenty-seven females provided blood samples in which p70S6K and phosphorylated p70S6K were analyzed. Chart review gathered biometric measurements, clinical phenotypes, and medication response. Questionnaires assessed anxiety, depression, autism traits, and mitochondrial dysfunction, to determine neuropsychiatric disease profiles. Univariate and multivariate statistical analyses were used to identify predictors of medication response.

**Results:** mTORC1 activity correlated highly with both classical biometrics (height, macrocephaly, pupil distance) and specific neuropsychiatric disease profiles (anxiety and autism). Across all cases, phosphorylated p70S6K was the best predictor for ketamine response, and also the best predictor for rapamycin response in a single instance.

**Conclusions:** The data illustrates the importance of mTORC1 activity in both observable body structure and medication response. This report suggests that a simple assay may allow cost-effective prediction of medication response.

## INTRODUCTION

Although formal biochemical analysis of aging pathways is a relatively novel academic discipline, medical modulation of nutrient sensing pathways in humans has been widespread since the approval of lithium in the United States in 1970. Lithium, a glycogen synthase kinase inhibitor, has proven efficacy not only in bipolar disorders [1] but also more recently in Alzheimer’s disease prophylaxis [2], and is known to extend *Drosophila* lifespan [3]. The combination of lithium with drugs that inhibit the mammalian target of rapamycin complex 1 (mTORC1) (by rapamycin) and mitogen-activated protein kinase (by trametinib) pathways, can, in *Drosophila*, increase lithium’s 11% lifespan extension to 48% [4].

The inexpensive generic drugs ketamine and rapamycin, that are efficacious in both chronic pain and depression [5, 6], modulate different nutrient sensing pathways. These medications have proximal impacts on the innate nitrogen sensing pathway in every cell in the body via mTORC1, although potentially in opposing directions. Ketamine blocks an extracellular glutamate sensor, the N-methyl-D-aspartate (NMDA) receptor, which is preferentially activated during high activity in nerve cells. The resulting decrease in voltage-gated calcium flux and mTORC2 downregulation may stabilize small G protein Rheb and activate mTORC1 [7, 8]. In contrast, rapamycin mimics amino acid restriction and subsequently downregulates glycolytic-dependent growth and inflammatory processes, resulting in mTORC1 inhibition [4].

Optimal population use of these inexpensive medications in patients requires identification of biomarkers which specify the dose requirements for each drug in each individual. Absent aggressive innovation in diagnostic technology, it is likely the prophylactic effects of lithium, ketamine, or rapamycin will be introduced into the community too late to protect hundreds of millions of individuals who will ultimately die from Alzheimer’s disease or widespread behavioral consequences of depression. An additional barrier is minimal commercial financial interest to address “the last mile problem,” educating the individual patient to take generic stigmatized medications lifelong with potential immediate side effect versus uncertain long-term benefit.

Treatment-resistant psychiatric patients, refractory to standard Food and Drug Administration (FDA) approved medications for depression, in contrast, actively demand novel treatments and diagnostics. Although high dimensional tissue assays, both genomic and metabolomic, are recently commercially available, in our experience, practical utility is limited by cost, convenience, excessive dimensionality of multiple findings, and uncertain applicability of observed peripheral somatic tissue pathophysiology to brain pathophysiology.

In response to patients’ unmet needs, we observed that recent developments in western blot technology and conceptual advances in biochemical pathways derived from yeast models potentially allow for cost-effective estimation of mTORC1 activity in the individual patient.

We hypothesized that biochemical activity in activated monocytes is a cost-effective estimate of baseline human mTORC1 gain of function. The cost ($45) and convenience of the acquisition of blood cells is trivial. Although blood is composed of a heterogeneous group of cells, from a biochemical perspective most cells in the blood have minimal mTORC1 activity [9]. The exception to this generalization may be monocytes; in mice models phosphorylation of p70 ribosomal S6 kinase (p70S6K), a downstream target of mTORC1, was found to be uniquely elevated in monocytes relative to neutrophil or lymphocyte subsets [9]. In physiological settings, Ly6hi “classical monocytes” with elevated Ki67+activity transition into terminally differentiated Ly6low tissue-resident monocytes during exposure to colony stimulating factor 1 (CSF-1) from damaged portions of the vascular endothelium [10, 11]. Pathological overgrowth of these transitioning cells may be observed in numerous varied disease phenotypes: systemic juvenile idiopathic arthritis [9], lithium-responsive bipolar disorder [1], and rapamycin treatment in amyotrophic lateral sclerosis [12]. Unfortunately, blood samples may be less than optimal as markedly higher expression of PI3K-AKT, glycolytic genes, and Hif1a are observed in wound “M1-like” macrophages at day 2 post injury compared with blood monocytes [13].

The objective of this study was to determine whether presumed macrophage mTORC1 activity, as measured by p70S6K phosphorylation, has implications in terms of identifying undiscovered patient phenotypes or indications for specific medications.

## RESULTS

### Clinical variables of the patients

Twenty-five patients and two first-degree relatives of patients were recruited for this study. Descriptive statistics of their clinical variables and scores on the questionnaires can be found in **Table 1**. The central tendency in this sample is consistent with membership in a community fee-for-service psychiatric clinic. The women are middle-aged, overweight, and with a normative height. They have a moderate degree of residual anxiety as inferred from the GAD-7. They score in the mid-range for autism traits on the AQ-10. The scores on the PHQ-9, an estimate of cognitive fatigue associated with fibromyalgia, are skewed towards substantial clinical disease. Based on these results, these patients are not normal, and not representative of a random community sample.

**Table 1.**
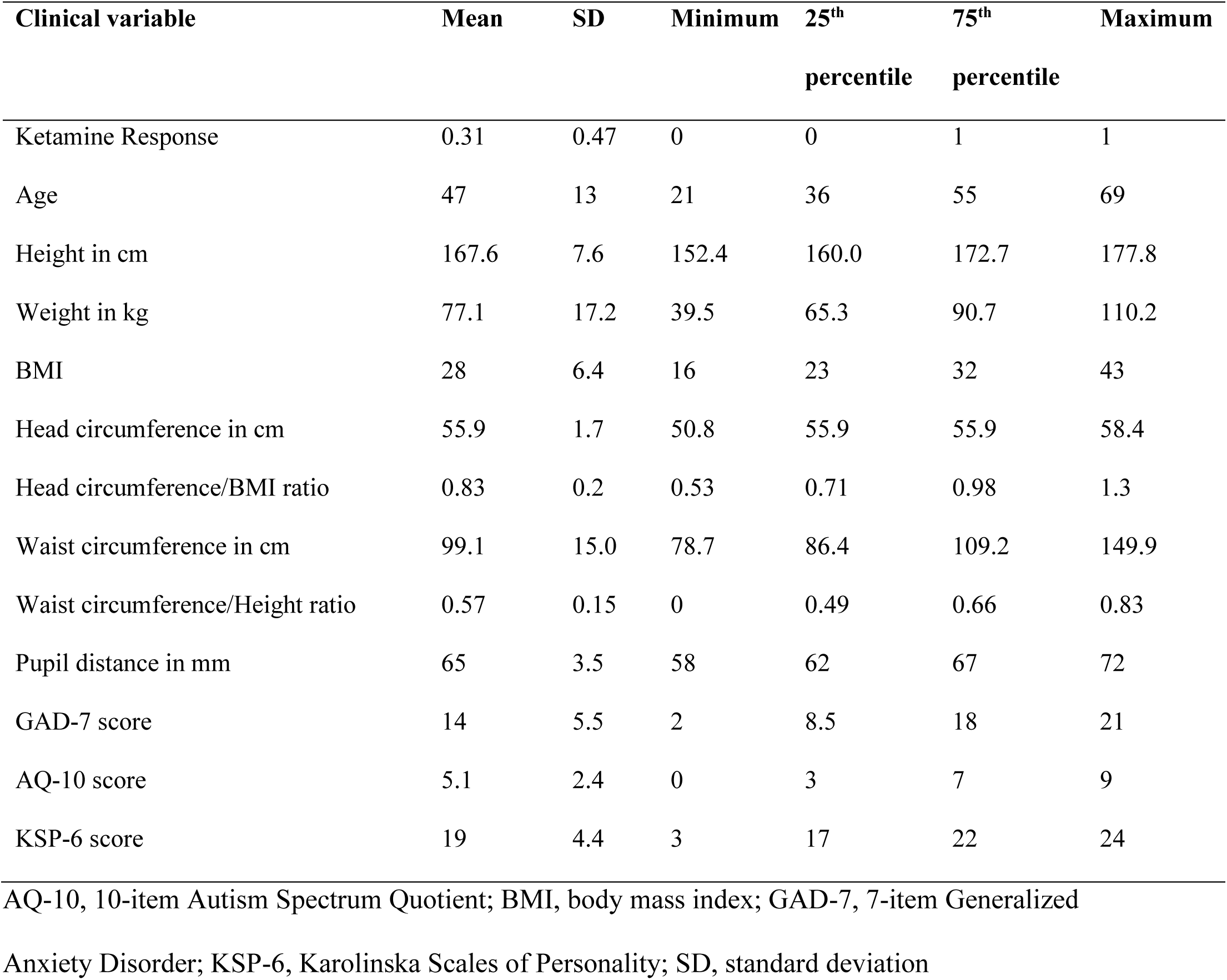
Summary statistics of the clinical features with percentiles.

| Clinical variable | Mean | SD | Minimum | 25<sup>th</sup><br>percentile | 75<sup>th</sup><br>percentile | Maximum |
| --- | --- | --- | --- | --- | --- | --- |
| Ketamine Response | 0.31 | 0.47 | 0 | 0 | 1 | 1 |
| Age | 47 | 13 | 21 | 36 | 55 | 69 |
| Height in cm | 167.6 | 7.6 | 152.4 | 160.0 | 172.7 | 177.8 |
| Weight in kg | 77.1 | 17.2 | 39.5 | 65.3 | 90.7 | 110.2 |
| BMI | 28 | 6.4 | 16 | 23 | 32 | 43 |
| Head circumference in cm | 55.9 | 1.7 | 50.8 | 55.9 | 55.9 | 58.4 |
| Head circumference/BMI ratio | 0.83 | 0.2 | 0.53 | 0.71 | 0.98 | 1.3 |
| Waist circumference in cm | 99.1 | 15.0 | 78.7 | 86.4 | 109.2 | 149.9 |
| Waist circumference/Height ratio | 0.57 | 0.15 | 0 | 0.49 | 0.66 | 0.83 |
| Pupil distance in mm | 65 | 3.5 | 58 | 62 | 67 | 72 |
| GAD-7 score | 14 | 5.5 | 2 | 8.5 | 18 | 21 |
| AQ-10 score | 5.1 | 2.4 | 0 | 3 | 7 | 9 |
| KSP-6 score | 19 | 4.4 | 3 | 17 | 22 | 24 |

AQ-10, 10-item Autism Spectrum Quotient; BMI, body mass index; GAD-7, 7-item Generalized
Anxiety Disorder; KSP-6, Karolinska Scales of Personality; SD, standard deviation

### Expression and phosphorylation of p70S6K

Phosphorylation of p70S6K, a downstream target of mTORC1, was measured as a proxy for mTORC1 activation in monocytes. The Western blot results are shown in **Supplementary** Figure 1. During analysis, difficulty in actin estimation occurred in 10 samples, limiting actin as a single control for variable aliquot sizing to 17 patients. However, the relative ratio of phosphorylated and unphosphorylated p70S6K could still be determined for all 27 samples (**Table 2**). Descriptive statistics of expression and phosphorylation of p70S6K of the samples are presented in **Table 3**.

**Table 2.**
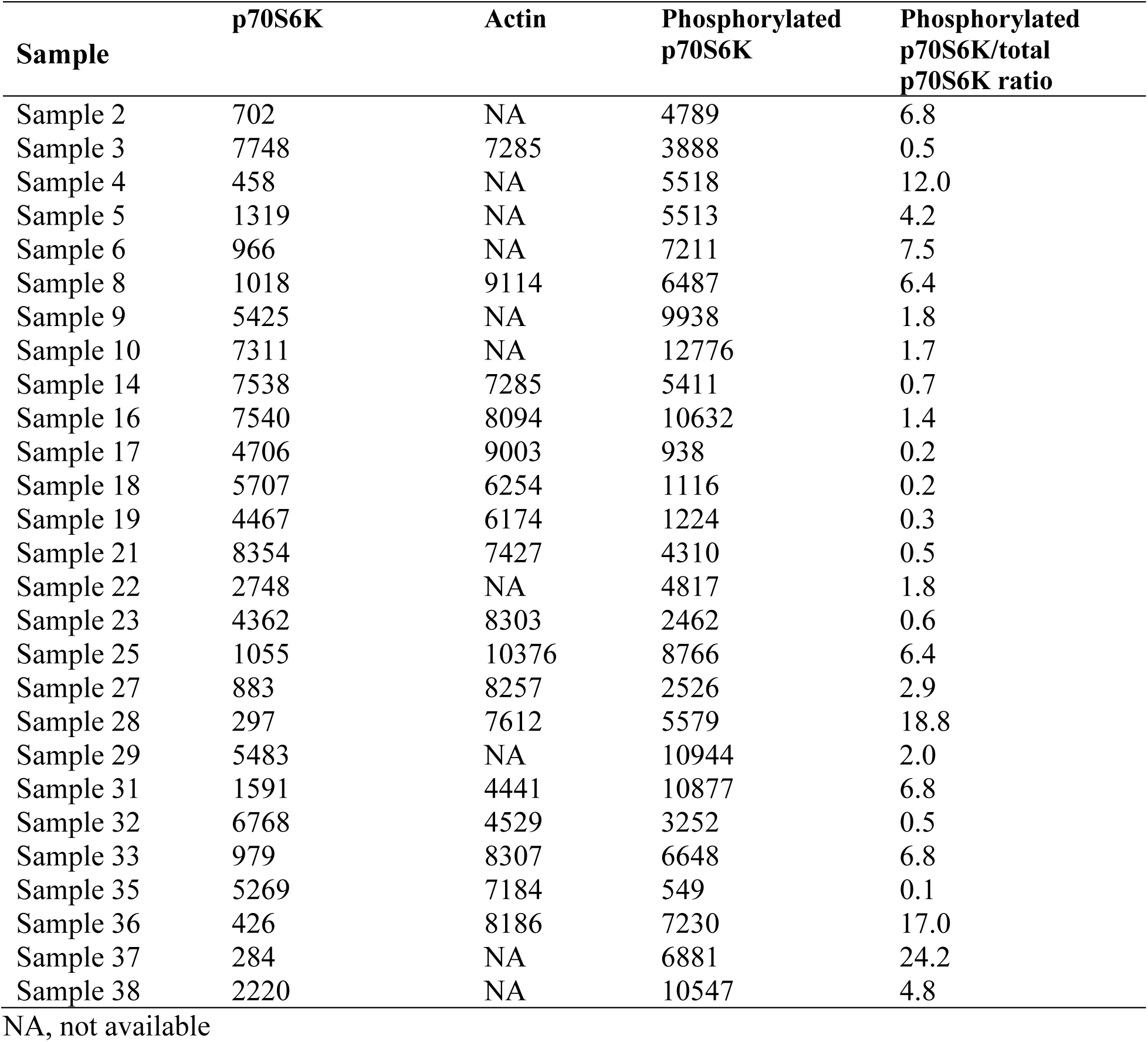
Western blot results of mTORC1 activity per sample.

**Table 3.**
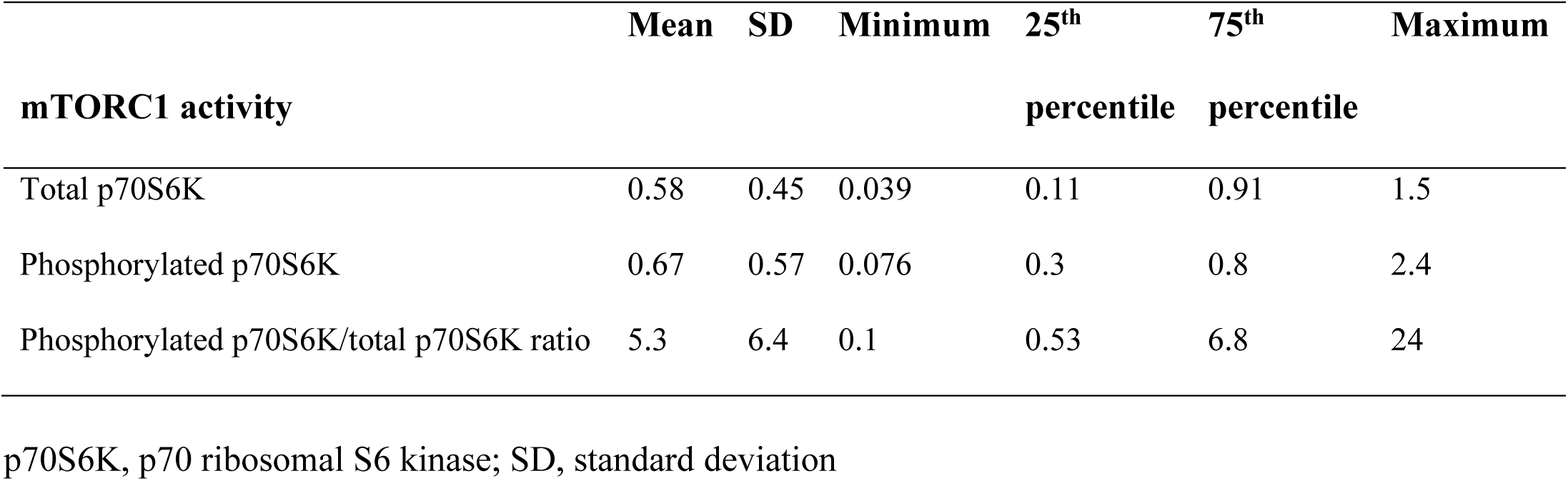
Overall mTORC1 activity.

### Clusters were identified based on multivariate analysis

Even within a highly selected population, women with anxiety and fibromyalgia, at most representative of 1-2% of the population, there was substantial within-group variability [14]. This within-group variability is the bane of clinical management of syndromic disease. Accordingly, we attempted to identify phenotype clusters potentially allowing for more targeted treatments.

Visual inspection of the correlation heatmap matrix (**Figure 1**) identified a few clusters. For example: One cluster was defined by the “antidiabetic” bobble-head frame (large head circumference relative to BMI), low p70S6K expression, and anxiety. Another cluster was defined by p70S6K expression, head circumference, and pupil distance.

**Figure 1.**
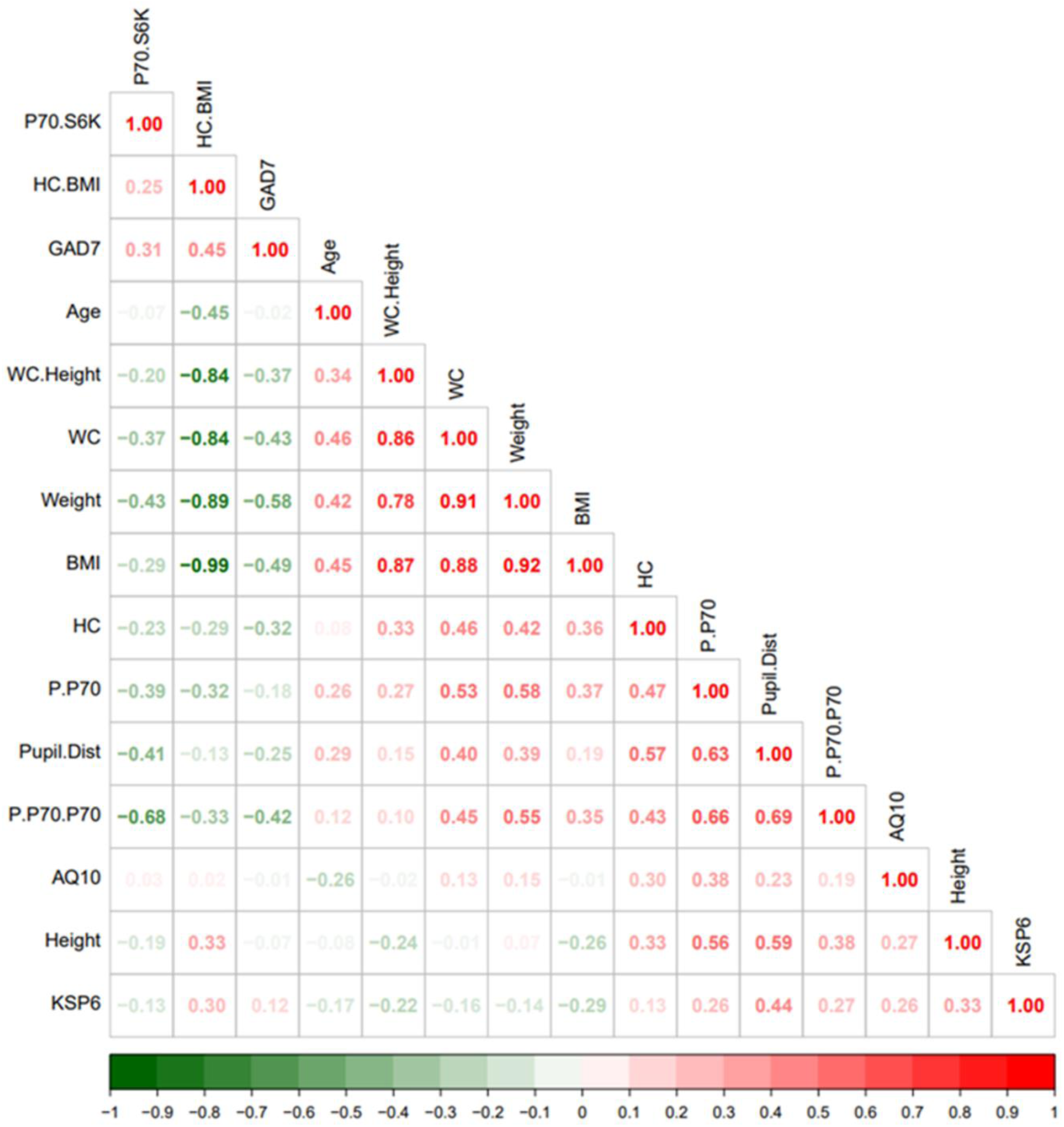
Heatmap visualization based on spearman rank correlation and hierarchical clustering. The legend bar shows positive (red) or negative (green) correlations between the various parameters.

### Principal component analysis identified two clusters

We also used another method of data compression, namely principal component analysis (PCA). PCA is extremely valuable as it allows for the identification of nearest neighbors defined by multiple variables rather than with a binary disease identification. PCA identified two principal components (PCs) accounting for more than 50% of the variance in a sample of 8 biometric variables and neuropsychiatric disease profiles (anxiety and autism) based on the GAD-7, KSP-6, and AQ-10 scores (**Supplementary** Figure 2). PC1 explained 36.5% and PC2 explained 18% variance, respectively.

### Phosphorylated p70S6K was the best predictor for ketamine response

We used ketamine response as an outcome variable (responded as “1” vs. not responded as “0”) and used the Random Forest method (as classification mode) to identify key clinical parameters that predict ketamine response. Random forest ranking using mean decrease in accuracy identified phosphorylated p70S6K, height, and ratio of head circumference/BMI as the top three predictive features (**Figure 2**). A list of all the parameters with their ranking based on Random Forest model can be found in **Supplementary Table 2.**

**Figure 2.**
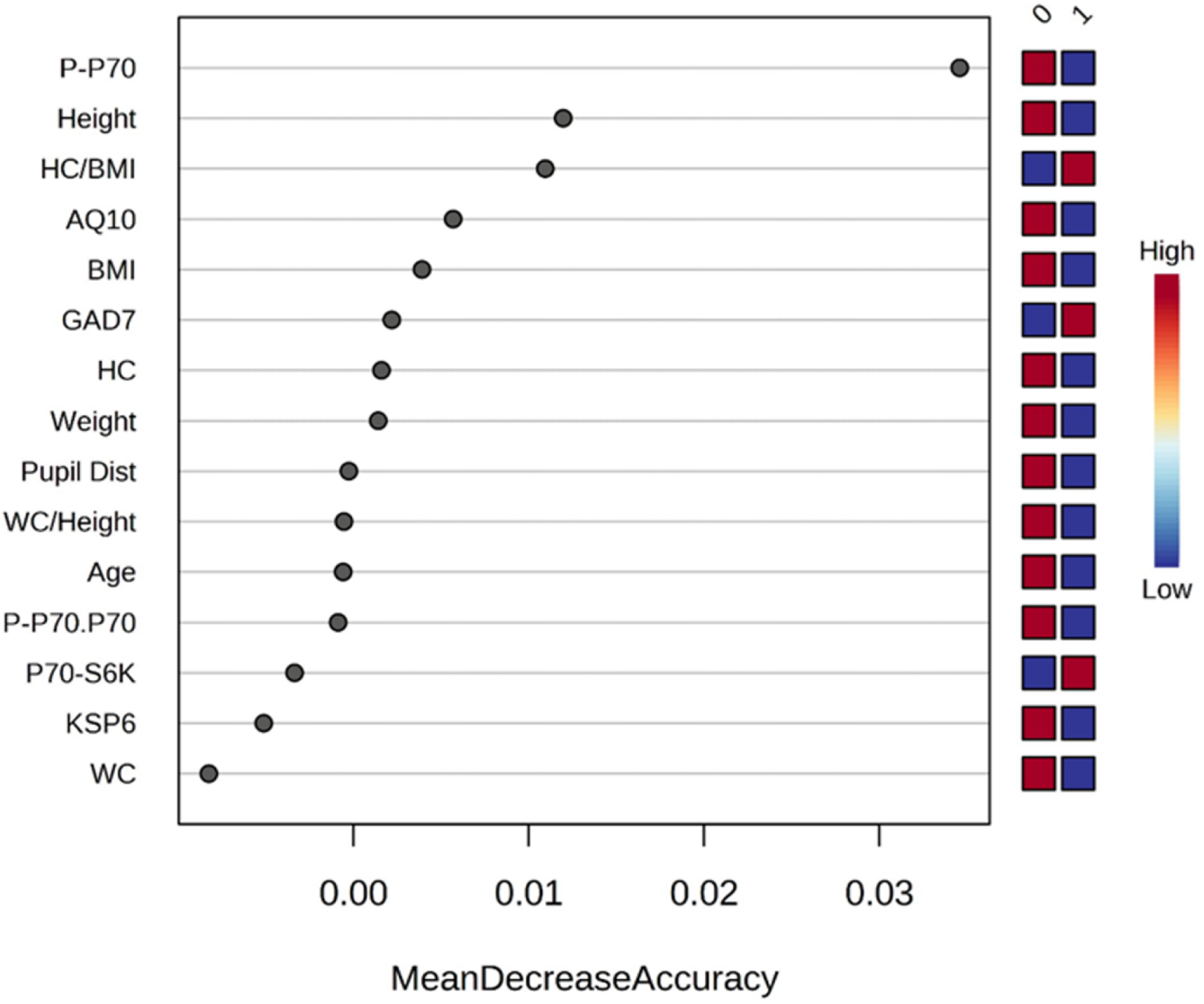
Rank of the clinical features that affect ketamine response based on the random forest algorithm. Ketamine response is indicated as “1” for responded and “0” for not responded. Corresponding up and downregulation of the parameters are indicated on the right side of the plot. AQ10, 10-item Autism Spectrum Quotient; BMI, body mass index; GAD7, 7-item Generalized Anxiety Disorder; KSP6, HC, head circumference; HC BMI, ratio head circumference/BMI ratio; Karolinska Scales of Personality; PCA, principal component analysis; Pupil Dist, pupil distance; P.P70, phosphorylated p70S6K; P.P70.P70, ratio of phosphorylated p70S6K/total p70S6K; WC, waist circumference; WC Height, ratio waist circumference/height.

**Figure 3:**
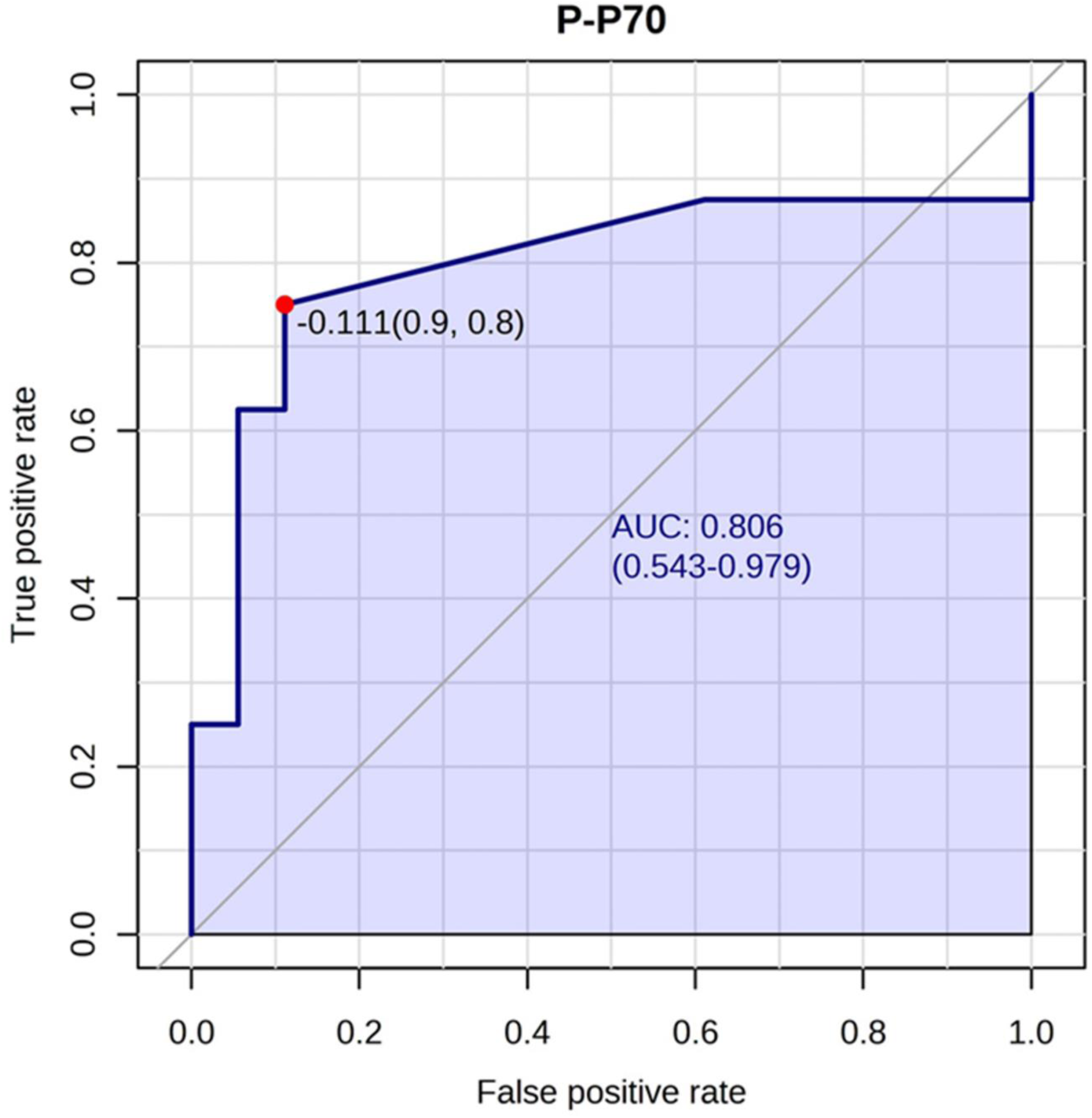
Phosphorylated p70S6K best predicted ketamine response. The phosphorylated p70S6K marker had an AUC of 0.80 (confidence interval: 0.54-0.97) for predicting ketamine response. An optimal cutoff is shown (indicated by the red dot) based on true positive and false positive rate.

We also performed univariate t-tests to estimate the parameters’ area under curve (AUC). A list of the AUCs can be found in **Supplementary Table 1**. The highest AUC was found for phosphorylated p70S6K (AUC 0.79), indicating this is a good predictor of ketamine response, and the lowest AUC was found for p70S6K expression (AUC 0.51).

### Rapamycin response correlated with phosphorylated p70S6K: Case example

A 30-year-old male patient with a chronic history of ulcerative colitis and idiopathic pancreatitis. At index presentation, he was occupationally disabled with crippling fatigue, intermittent hospitalizations for intractable vomiting and pain, and severe cognitive impairment.

Psychostimulant use was palliative in contrast to previous antidepressants and standard immunosuppressive agents. In the course of standard clinical discussion regarding the use of rapamycin in treatment-resistant lupus and subsequent Treg induction, his mother volunteered to provide a tissue sample consistent with known heritability of autoimmunity. Subsequent evaluation revealed the largest amount of phosphorylated p70S6K in this clinical sample, suggesting elevated Bayesian probability for rapamycin indication. The patient consented to a 1 mg daily rapamycin trial under Washington State Right-to-Try statutes with a first-degree relative present (See **Informed Consent Form** in **Supplemental Data**). After a four-week trial, the patient reported complete cessation of fatigue with an exercise tolerance of 12 hours, spontaneous cessation of lifelong 1.5 pack daily nicotine dependence with one week, and minimal self-imposed dietary restriction associated with residual disease (1/30 days).

Patient statement: “*After continued use of rapamycin, I have found several noticeable changes in my emotional and physical states. My dependency on nicotine is no longer existent and I have fully quit without implementing any secondary forms of cessation. My severe ulcerative colitis, Crohn’s, and pancreatitis flare-ups have become increasingly rare. On the off chance they do happen, my recovery time is incredibly fast. I have been able to go from a very restrictive diet to prevent them to now being able to have a much more varied and tolerable diet. My energy levels have become extremely high and I am able to work as long and hard at whatever task I’m working on. This includes minimal fatigue and recovery time afterwards as well. Lastly, my overall mental health has become very much improved. My depression has become very minimal with very few bad days. I’ve been much happier to the point where I have actively started creating a social life. I haven’t had the desire, energy, or positivity to do this in several years*.”

## DISCUSSION

Our data suggests that human variability of mTORC1 gain of function observed during the differentiation of stem-like monocytes into vascular tissue-resident macrophages correlates with physical size, subsets of neuropsychiatric disease, and clinical ketamine or rapamycin response.

The major strength of this study is the absence of commercial interests. Funding for this study ultimately derived from patients with treatment-refractory disease with a potentially life-threatening course. All medications of interest studied were generic, historically underutilized, and relevant to their probable benefit absent commercial marketing. No professional conflicts of interest regarding patents, pending grant applications, or publication prominence required for professional advancement were present. In the absence of commercial conflicts and subsequent required regulatory delays, the latency between project design and successful clinical application was short, six months. This latency is entirely consistent with the intent of recent regulatory developments in Federal and Washington state Right-to-Try laws [15].

This study has several limitations. First, this study involved only female patients and cannot be extrapolated to males. So, a study in a male cohort is required given elevated bimodal expression of synaptic disease in males (overgrowth in autism vs. undergrowth in schizophrenia) [16, 17], decreased relative risk of hyperinflammatory manifestations (somatic autoimmune disease and central nervous system mood disorders) [18], and the highly variable differences in response to life-extending interventions between sexes, such as reduced IGF-1 or mTOR signaling, observed in mice [19]. Second, to confirm the proposed tentative model, isolation of monocyte subpopulations from the blood using cell sorting is needed. Third, non-linear distributions of mTORC1 activity require further investigations using alternative readouts e.g., 4E-binding protein 1 (4EBP1) or AKT subtypes, which might allow for a more robust biochemical gradient when estimating medication selection and dose. Fourth, the results of multiple non-preregistered statistical comparisons within a restricted clinical sample require confirmation in a study with a larger number of patients before applying the observed effect sizes to broader populations.

The strong association between a fixed physical structure (i.e., head size and pupil distance) and dynamic macrophage differentiation is consistent with known mTORC1 effects on macrocephaly [20]. This finding provides face validity to the concept that systemic mTORC1 gain of function can be estimated easily, albeit incompletely, with simple physical measurements and crude blood analysis. Age, exercise, and diet-related hypertrophic lipodystrophy manifest as increased waist circumference and BMI and may rely on different proximal upstream activators of mTORC1 activated later in development than in infancy and adolescence.

The miscellaneous associations between multiple neuropsychiatric syndromes (anxiety and autism) and mTORC1 activity in blood monocytes direct attention to the structural role of tissue-resident macrophages in the brain, the microglia. In the brain, are physical tension and psychological tension the same? The brain is a closed space due to the inflexibility of the skull. All learning is a zero-sum game. Therefore, an augmented synaptic trace associated with a new experience requires the corresponding phagocytosis of a less preferred synapse. The extracellular matrix, in turn, must be digested to avoid “tension” as different synaptic “eigenvalues” embedded in brain activity “eigenvector” grow in size with time and distort the morphology of connecting tissue or present risk of local excitotoxicity [21]. This conceptual framework suggests that different densities of the extracellular matrix and embedded synaptic structure may manifest as dimensional behavioral traits in humans.

The association of mTORC1 activity with ketamine response is consistent with the literature. Reductions in serum TNF-α were found to correlate with clinical antidepressant response after intravenous ketamine treatment; these changes were observed within 40 minutes [22]. Recent work in invasive vs. non-invasive ovarian cancer revealed that high levels of tumor-derived N-acetyl aspartate competitively inhibited the NMDA receptor on M1 tissue-resident macrophages and induced M2 polarization [23]. The resultant biochemical profile within the proliferative MKI67+ M2 macrophage (GLS1 upregulation, SLCA5/SLCA7 upregulation) strongly corresponded to the biochemical phenotype observed in the whole blood, inferred monocyte fraction, in human lithium responders [1]. These converging observations suggested the possibility that the M2 homeostatic microglia population (CD163+) in the brain of depressed patients is of inadequate size to allow for adequate synaptic pruning or trophic support [8]. The upregulation of BCL-2 in lithium responders [1] could potentially suggest protection against premature senescence of the long-lived microglia stem cell population, which migrates into the brain from the yolk sac at week 4 of embryonic development.

The association of mTORC1 activity with rapamycin response is consistent with the literature, specifically rapamycin’s efficacy in treating tuberous sclerosis [24]. Rapamycin also has documented efficacy in a host of polygenetic autoimmune disorders, specifically in the prototype disorder lupus [25] and the case-specific disorder ulcerative colitis [26]. Its proximal mechanism of action may relate to biasing differentiation of naive T-cells towards the FoxP3+ T-regulatory and CD8+CD45R0+ memory subtypes dependent on fatty acid oxidation and reducing interleukin (IL)-4 and IL-17 production by CD3+CD4-CD8-double negative T-cells.

The observed disjunction between relative phosphorylated p70S6K and absolute p70S6K abundance correlations with clinical phenotypes suggested careful attention to the different physiological drivers of metabolic reprogramming in macrophages. The total amount of available p70S6K and its active phosphorylated form may only partially control macrophage activity. Note that phosphoinositide 3-kinase (PI3K)-γ inhibition alone promoted the “M1-phenotype (IL-12 excretion)” relative to the “M2-phenotype (IL-10 excretion)” in the context of lipopolysaccharide (LPS) stimulation without affecting p70S6K activity [27]. Even within the “M2-phenotype” in cultured microglial cells, PI3K/TORC1 independent (IL-4) and dependent (ATP/purine) extracellular receptor activation pathways appeared to be present [28].

In conclusion, recent conceptual, technological, and regulatory developments may allow for the rapid development of a simple blood assay supporting both widespread and yet targeted use of generic medications known to extend both healthspan and lifespan. This blood assay attempts to capture a snapshot of “stem-like” monocytes that will transform into tissue-resident macrophages. The resulting image of innate mTORC1 and mTORC2 activation may provide insights into the management of neuropsychiatric disease. The current study provides a proof of concept that requires replication in larger cohorts and using isolated monocyte subpopulations.

## MATERIALS AND METHODS

### Patients and samples

Patients or first-degree relatives of patients with treatment-refractory disease were recruited from a community fee-for-service psychiatric practice in the course of chronic medication and case management between January and March of 2023. Patient selection was informal, largely in the background of discussions of synaptic undergrowth and overgrowth syndrome associated with autism/pain/mood disorders or modulation of aging-related biochemical pathways. Enrollment was entirely driven by patient interest, absent available pilot data suggesting immediate efficacy or application.

We assessed height, weight, head circumference, pupil distance, current medication, and age. Self-administered questionnaires were used to assess the presence of various disorders. Depression was assessed with the 9-item Patient Health Questionnaire (PHQ-9) [29], anxiety with the 7-item Generalized Anxiety Disorder (GAD-7) questionnaire [30], presence of autism traits with the 10-item Autism Spectrum Quotient (AQ-10) questionnaire [31], and mitochondrial dysfunction was assessed with the 6-item Karolinska Scales of Personality (KSP-6) questionnaire [32]. Retrospective chart review identified individuals who had or were receiving benefit from self-administered ketamine or rapamycin therapy; response was scored as “1” for responded and “0” for not responded. The response amplitude was not estimated; patient sustained use was taken as a marker of efficacy.

Fasting blood samples were collected in EDTA tubes between 8 AM and 12 PM within 2 weeks of clinical evaluation. Blood samples were centrifuged for 30 minutes at 15,000 rpm and an aliquot of cells was visually extracted. The samples were frozen on dry ice immediately for later examination.

### Informed consent

The retrospective case review was exempt from Institutional Review Board approval under Category 2 of the Basic Health and Human Services Policy for Protection of Human Research Subjects Subpart A Section 46.101 [33]. All participants signed informed consent forms (see **Supplemental Data**) consistent with Washington State Right-to-Try statutes Chapter 69.77 RCW [15]. Elements of the Right-to-Try statutes include acknowledgment of life-threatening disease in the patient or relative, absence of available investigational treatments or diagnostics, and assumption of personal financial responsibility for proposed interventions.

### Western blot analysis

To examine the expression and phosphorylation of p70S6 kinase in each sample, western blot analysis was carried out. To extract the proteins, 5 µl of each cell sample was mixed with 300 µl radioimmunoprecipitation assay (RIPA) buffer (25 mM Tris-HCl pH 7.4, 150 mM NaCl, 1% TritonX-100, 1% sodium deoxycholate, 0.1% sodium dodecyl sulfate (SDS), 1 mM ethylenediaminetetraacetic acid (EDTA), 5% glycerol, 1% 100 X Protease Inhibitor Cocktail) and kept on ice for 30 min. Samples were homogenized with a Branson SFX150 Ultrasonic Cell Disruptor (Emerson, UK) at amplitude setting: 40%; sonication pulse rate: 10 seconds ON, 10 seconds OFF, for 30 seconds on ice. Homogenized samples were kept on ice for 15 minutes and then centrifuged at 13,000 rpm for 15 min at 4°C. The supernatants containing the soluble protein were collected and transferred to new tubes. Proteins were separated on 10% Bis-Tris SDS gel. After transferring the proteins from the gel to a polyvinylidene difluoride membrane, subsequent treatments (blocking, antibody treatment) were performed using 5% bovine serum albumin (BSA) solution prepared in 1X Tris-buffered Saline, 0.1% Tween20. Antibodies against non-phosphorylated p70 S6 kinase (Cell Signaling, catalog no:2708), or phosphorylated p70 S6 kinase (Cell Signaling, catalog no: 9234) were used, as well as against actin (Thermo Fisher, catalog no: MA1-744) as an internal (protein loading) control. Antibody dilutions were prepared according to the manufacturer’s protocol.

### Analysis of Western blot results for measuring mTORC1 activity

Relative optical densities of target proteins were measured with ImageJ (Version 1.53t) software, and normalized to those of β-actin, when available. The β-actin bands for some samples were oversaturated during imaging, so these were instead normalized as a ratio of P-p70S6K/p70S6K.

Visual inspection of raw data revealed non-Gaussian distribution and/or outliers in many variables, suggesting Spearman correlation analysis. All clinical variables were then examined in a correlation matrix without preregistered hypothesis testing.

### Univariate and multivariate statistical analysis

Spearman correlation or association analysis was performed to understand the linking of the multiple clinical parameters. A univariate t-test was performed for Ketamine response as outcome variable. P value <0.05 was considered for statistical significance.

Principal component analysis (PCA) was used to decrease the dimensionality of a given dataset while retaining the utmost significant information. This was achieved through the identification of a novel collection of variables, referred to as the principal components (PCs), which exhibit no correlation and effectively capture the highest amount of variance present in the dataset. A total of 21 missing values in the features were detected in the twenty-seven patients. The missing values were imputed using median values, and the data was normalized using auto scaled (Z-transformation).

We have used R (https://www.r-project.org/) and metabolanalyst (https://www.metaboanalyst.ca/) tools for data analysis.

Random forest analysis, a type of ensemble supervised machine learning, was used to make the results more stable by combining the predictions of several decision trees [34, 35]. Random forest analysis was used for both classification and regression. We ranked (in decreasing order) the features based on mean decrease in accuracy.

## ABBREVIATIONS

AUC: Area under the curve
AQ-10: 10-item Autism Spectrum Quotient
BMI: Body mass index
BSA: Bovine serum albumin
CI: Confidence interval
EDTA: Ethylenediaminetetraacetic acid
FDA: Food and drug administration
GAD-7: 7-item Generalized Anxiety Disorder
IL: Interleukin
KSP-6: 6-item Karolinska Scales of Personality
LiTMUS: Lithium Treatment-Moderate dose Use Study
mTORC1: mammalian target of rapamycin complex 1
NMDA: N-methyl-D-aspartate
PC: Principal component
PCA: Principal component analysis
PHQ-9: 9-item Patient Health Questionnaire
RIPA: radioimmunoprecipitation assay
SD: Standard deviation
SDS: Sodium dodecyl sulfate
p70S6K: 70 ribosomal S6 kinase
UMAP: Uniform manifold approximation and projection
4EBP1: 4E-binding protein 1

## DISCLOSURES

### Author Contributions

K.B.: Study concept, data acquisition, data management, manuscript revision; N.O.: Data acquisition; A.K.: Data acquisition, revision of manuscript; A.A.: Statistical analysis, co-writing first draft of the manuscript, revision of the manuscript; J.B.: Study concept, plan of analysis, literature review, manuscript write-up and revisions.

## Acknowledgements

We thank Alexander Medenhall, PhD for arranging the collaboration, and Ayush Sharma and Alessandro Bitto, PhD for Western Blot analyses.

## Ethical Statement

The retrospective case review is exempt from Institutional Review Board approval under Category 2 of the Basic Health and Human Services Policy for Protection of Human Research Subjects Subpart A Section 46.101.

## Consent

All participants gave informed consent for the study by signing an informed consent form.

