## Supplemental Materials for "mTORC1 Activation In Presumed Classical Monocytes: Observed Correlates With Human Size Variation and Neuropsychiatric Disease"

### SUPPLEMENTAL DATA

**Informed Consent Form**

RAPAMYCIN RIGHT-TO-TRY

RAPAMYCIN FOR PREVENTION AGE-RELATED PSYCHIATRIC DECLINE

RIGHT TO TRY IN WASHINGTON STATE CONSENT FORM

1/17/21

"Section 1: The legislature finds that the process for approval of investigational drugs, biological products, and devices in the United States protects future patients from premature, ineffective and unsafe medications and treatments over time, but the process often takes many years. Patients who have a terminal illness do not have the luxury of waiting until an investigational drug, biological product or device receives final approval from the Unites States food and drug administration. The legislature further finds that patients who have a terminal illness should be to pursue the preservation of their own lives…."

1. _____ I agree that I have a serious or immediately life threatening disease (reasonable likelihood of death within 6 months OR premature death is likely without treatment)
2. _____I understand that I have failed a reasonable subset of established treatments for my illness and/or unwilling to pursue high risk heroic treatments
3. _____I understand that there are no major therapeutic advances available in the pipeline in terms of local clinical trials.

4. ____I understand in the WORST CASE rapamycin could produce immunosuppression, damage to the lining of the mouth and stomach, worsen underlying diabetes, or fluid leak in the lungs or limbs. Many of these conditions could produce premature death. There may also be side effects of this drug that have not yet been discovered.
5. ___I understand in the BEST CASE rapamycin may slow the expected progression of my illness with age. There is additional animal and human data suggesting the potential for recovery of function across a wide variety of illnesses. Your individual probability of response cannot be estimated with current technology.
6. ___I understand that my insurance carrier may not be liable for treatment costs or any harm caused by rapamycin

**Supplementary Table 1 Clinical parameters and corresponding AUC values for ketamine response prediction**

| Clinical parameter | AUC (95% CI) |
|---|---|
| Phosphorylated p70S6K | 0.80 (0.53–0.97) |
| Ratio phosphorylated p70S6K/total p70S6K | 0.78 (0.59–0.94) |
| Weight | 0.76 (0.54–0.93) |
| Head circumference | 0.74 (0.50–0.93) |
| Pupil Distance | 0.74 (0.50–0.93) |
| Waist circumference | 0.74 (0.48–0.93) |
| AQ-10 score | 0.73 (0.51–0.91) |
| BMI | 0.72 (0.52–0.91) |
| Ratio head circumference/BMI | 0.72 (0.48–0.90) |
| GAD-7 score | 0.67 (0.47–0.91) |
| Height | 0.66 (0.48–0.91) |
| Ratio waist circumference/height | 0.64 (0.45–0.91) |
| KSP-6 score | 0.61 (0.45–0.90) |
| Age | 0.58 (0.42–0.90) |
| p70S6K expression | 0.51 (0.41–0.90) |

AUC, area under the curve; AQ-10, 10-item Autism Spectrum Quotient; BMI, body mass index; CI, confidence interval; GAD-7, 7-item Generalized Anxiety Disorder; KSP-6, Karolinska Scales of Personality

**Supplementary Table 2 Ranking of clinical parameters based on random forest modelling**

| Clinical parameter | Mean Decrease Accuracy |
|---|---|
| Phosphorylated p70S6K | 0.034579 |
| Height | 0.011966 |
| Ratio head circumference/BMI | 0.010941 |
| AQ-10 score | 0.00569 |
| BMI | 0.003914 |
| GAD-7 score | 0.002192 |
| Head circumference | 0.001602 |
| Weight | 0.001418 |
| Pupil distance | -0.00026 |
| Ratio waist circumference/height | -0.00054 |
| Age | -0.00058 |
| Ratio Phosphorylated p70S6K/total p70S6K | -0.00087 |
| p70S6K expression | -0.00336 |
| KSP-6 score | -0.00511 |
| Waist circumference | -0.00824 |

**Supplementary Figure 1 Western blots to assay phosphorylated p70S6K/total p70S6K ratio**

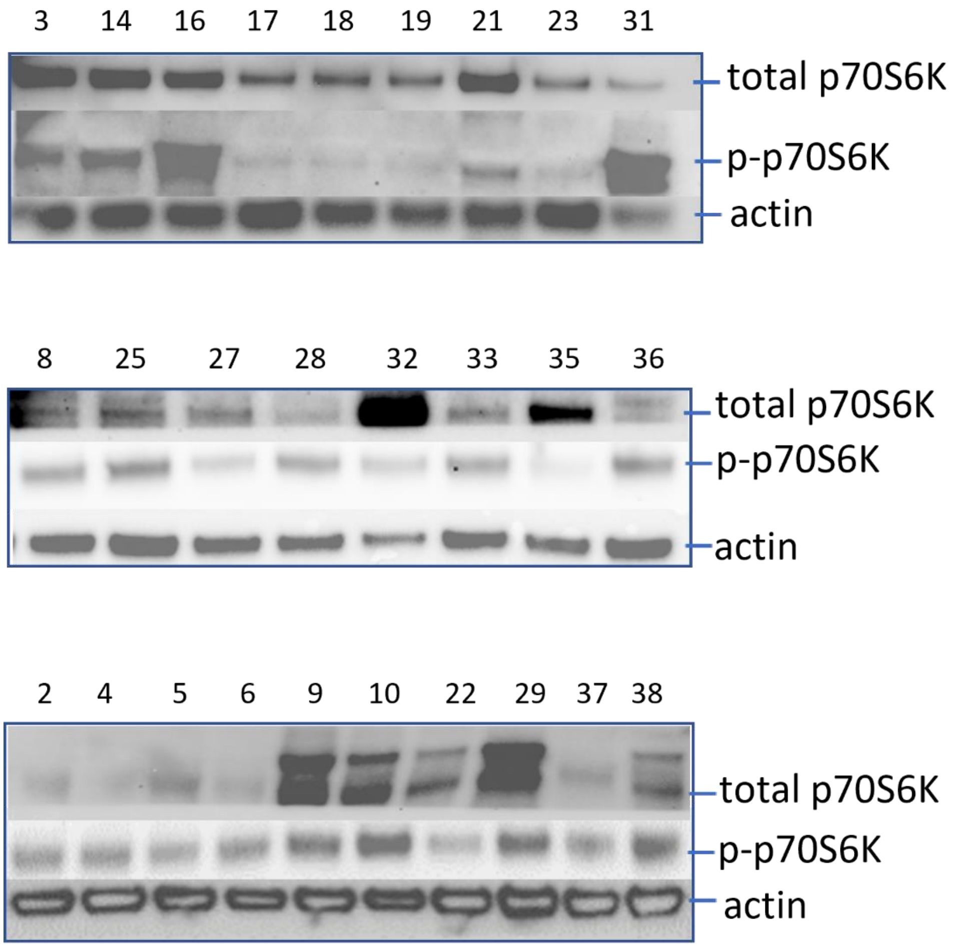


Western blot of p70S6K, phosphorylated p70S6K, and actin to assay the phosphorylated p70S6K/total p70S6K ratio. The actin quantity could not be estimated for the 10 samples in the bottom panel, limiting actin as control for aliquot sizing to 17 patients. Each number refers to a patient.

p70S6K, p70 ribosomal S6 kinase; p-p70S6K, phosphorylated p70S6K

**Supplementary Figure 2 PCA analysis on the entire dataset**

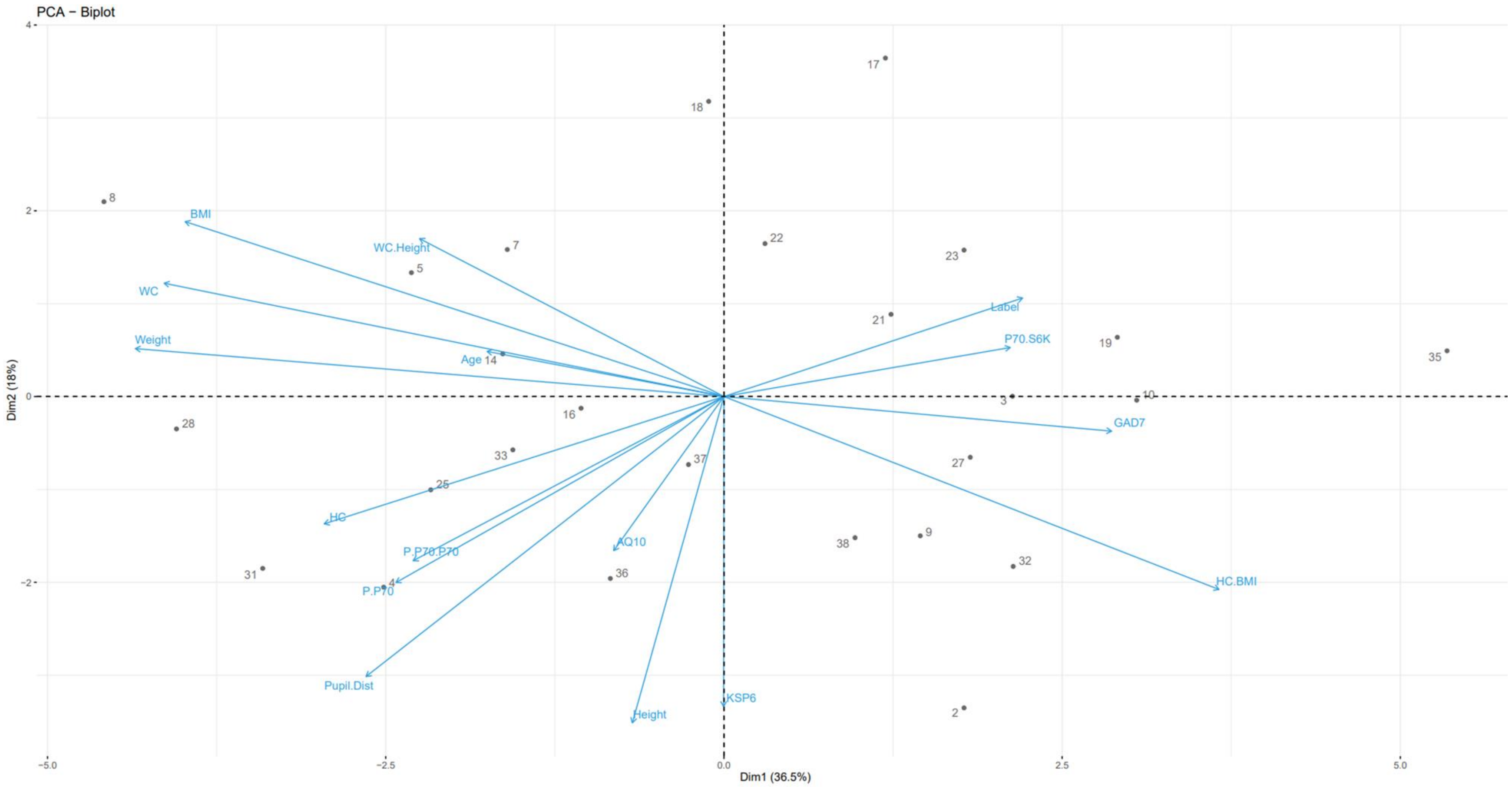


Each dot represents a patient. Dimension 1 (Dim1) explains 36.5% of the variance and dimension 2 (Dim2) explains 18% of the variance. "Label" indicates the ketamine response.

AQ10, 10-item Autism Spectrum Quotient; BMI, body mass index; GAD7, 7-item Generalized Anxiety Disorder; KSP6, HC, head circumference; HC BMI, ratio head circumference/BMI ratio; Karolinska Scales of Personality; PCA, principal component analysis; Pupil Dist, pupil distance; P.P70, phosphorylated p70S6K; P.P70.P70, ratio of phosphorylated p70S6K/total p70S6K; WC, waist circumference; WC Height, ratio waist circumference/height.
